# Detection of pathogenic splicing events from RNA-sequencing data using dasper

**DOI:** 10.1101/2021.03.29.437534

**Authors:** David Zhang, Regina H. Reynolds, Sonia Garcia-Ruiz, Emil K Gustavsson, Sid Sethi, Sara Aguti, Ines A. Barbosa, Jack J. Collier, Henry Houlden, Robert McFarland, Francesco Muntoni, Monika Oláhová, Joanna Poulton, Michael Simpson, Robert D.S. Pitceathly, Robert W. Taylor, Haiyan Zhou, Charu Deshpande, Juan A. Botia, Leonardo Collado-Torres, Mina Ryten

## Abstract

Although next-generation sequencing technologies have accelerated the discovery of novel gene-to-disease associations, many patients with suspected Mendelian diseases still leave the clinic without a genetic diagnosis. An estimated one third of these patients will have disorders caused by mutations impacting splicing. RNA-sequencing has been shown to be a promising diagnostic tool, however few methods have been developed to integrate RNA-sequencing data into the diagnostic pipeline. Here, we introduce *dasper*, an R/Bioconductor package that improves upon existing tools for detecting aberrant splicing by using machine learning to incorporate disruptions in exon-exon junction counts as well as coverage. *dasper* is designed for diagnostics, providing a rank-based report of how aberrant each splicing event looks, as well as including visualization functionality to facilitate interpretation. We validate *dasper* using 16 patient-derived fibroblast cell lines harbouring pathogenic variants known to impact splicing. We find that *dasper* is able to detect pathogenic splicing events with greater accuracy than existing *LeafCutterMD* or z-score approaches. Furthermore, by only applying a broad OMIM gene filter (without any variant-level filters), *dasper* is able to detect pathogenic splicing events within the top 10 most aberrant identified for each patient. Since using publicly available control data minimises costs associated with incorporating RNA-sequencing into diagnostic pipelines, we also investigate the use of 504 GTEx fibroblast samples as controls. We find that *dasper* leverages publicly available data effectively, ranking pathogenic splicing events in the top 25. Thus, we believe *dasper* can increase diagnostic yield for a pathogenic splicing variants and enable the efficient implementation of RNA-sequencing for diagnostics in clinical laboratories.

## Introduction

Next-generation sequencing has greatly accelerated the discovery of novel gene-to-disease associations^1,2^. As a result, whole exome sequencing (WES) and more recently, whole genome sequencing (WGS) are increasingly incorporated into the genetic diagnostic routine. However, it is estimated that the success rate of such DNA-sequencing approaches in Mendelian diseases is plateauing at 35-50%^3,4^. To an extent, this is due to the challenges of interpreting genetic variation beyond those that alter protein sequence or DNA structure^5,6^. In particular, non-coding regulatory variants remain difficult to assess and are more likely to be classified as variants of unknown significance (VUS), as compared to coding variants for which more analytic approaches exist^7^. Pathogenic variants that impact splicing are one class of non-coding variation, which are likely to account for a significant proportion of unsolved cases^8^. The splicing machinery is tightly regulated by numerous cis and trans signals; this complexity is crucial for generating transcript and phenotypic diversity, but also increases the likelihood that genetic variation will disrupt splicing^9,10^. In fact, variants distributed in non-coding regions of the genome disproportionately affect splicing, often through disruptions to intronic splicing enhancers, silencers or recognition sequences^11^. Furthermore, aberrant splicing has been shown to be a primary cause of rare diseases, with an estimated one third of pathogenic variants impacting splicing^12,13^.

Given the prevalence of unsolved rare disease patients with putative genetic causes through disruptions to splicing, there has been growing interest in the application of RNA-sequencing (RNA-seq) for diagnostics to directly measure transcriptome-wide splicing^14^. Using RNA-seq, we can obtain a functional readout of splicing levels, gene expression and allele-specific expression (ASE) in patients relative to unaffected controls. This enables the discovery of aberrant molecular products, which can be used to resolve the list of candidate genes and variants identified through WGS/WES to an actionable number. Aberrant RNA-level events discovered in this way can be used to re-prioritise VUS, leading to assignment of pathogenicity. Previous publications have demonstrated the promising utility of RNA-seq for diagnostics, with success rates ranging from 7.5-21% for patients with no candidate genes after WES and/or WGS^15–18^. In principle, information on splicing, gene expression and ASE obtained from RNA-seq all have diagnostic potential. However, in practice the majority of genetic diagnoses made through RNA-seq have involved detection of aberrant splicing^15,16^.

Since the first systematic application of RNA-seq for diagnostics by Cummings and colleagues in 2017, there has been growing interest in developing methods to detect pathogenic RNA events in rare disease patients^19–22^. Although numerous tools exist to perform differential splicing analysis, almost all are designed to identify global transcriptional differences between moderate-to-large case-control cohorts^23,24^. Few are specialised for genetic diagnosis, where success relies on distinguishing a pathogenic splicing event in a single patient (N of 1). Improvements to the methodologies to detect pathogenic splicing events will relieve clinical scientists of the requirement for manual curation, permitting the wider implementation of RNA-seq-based approaches within accredited diagnostic laboratories and increasing diagnostic success.

Here, we introduce *dasper*, a method which integrates disruptions in both exon-exon junction and base pair level coverage data through machine learning to detect aberrant splicing events in patient samples. We find that *dasper* detects pathogenic splicing events with greater accuracy than existing methods. After applying an OMIM-morbid gene filter, *dasper* is able to rank true pathogenic splicing events in the top 10 most aberrant splicing events. Furthermore, *dasper* is designed with diagnostic applications in mind and includes functionality to visualize candidate genes in the form of sashimi plots for manual inspection (Figure 1). Finally, we demonstrate that *dasper* is able to effectively leverage publicly-available control RNA-seq datasets, making RNA-seq a more cost-effective, standardized solution for diagnostics. *dasper* is released as an R package on Bioconductor (http://www.bioconductor.org/packages/dasper) and we believe that its use will improve the detection of pathogenic splicing events and, ultimately, the diagnostic yield for rare disease patients.

**Figure 1.**
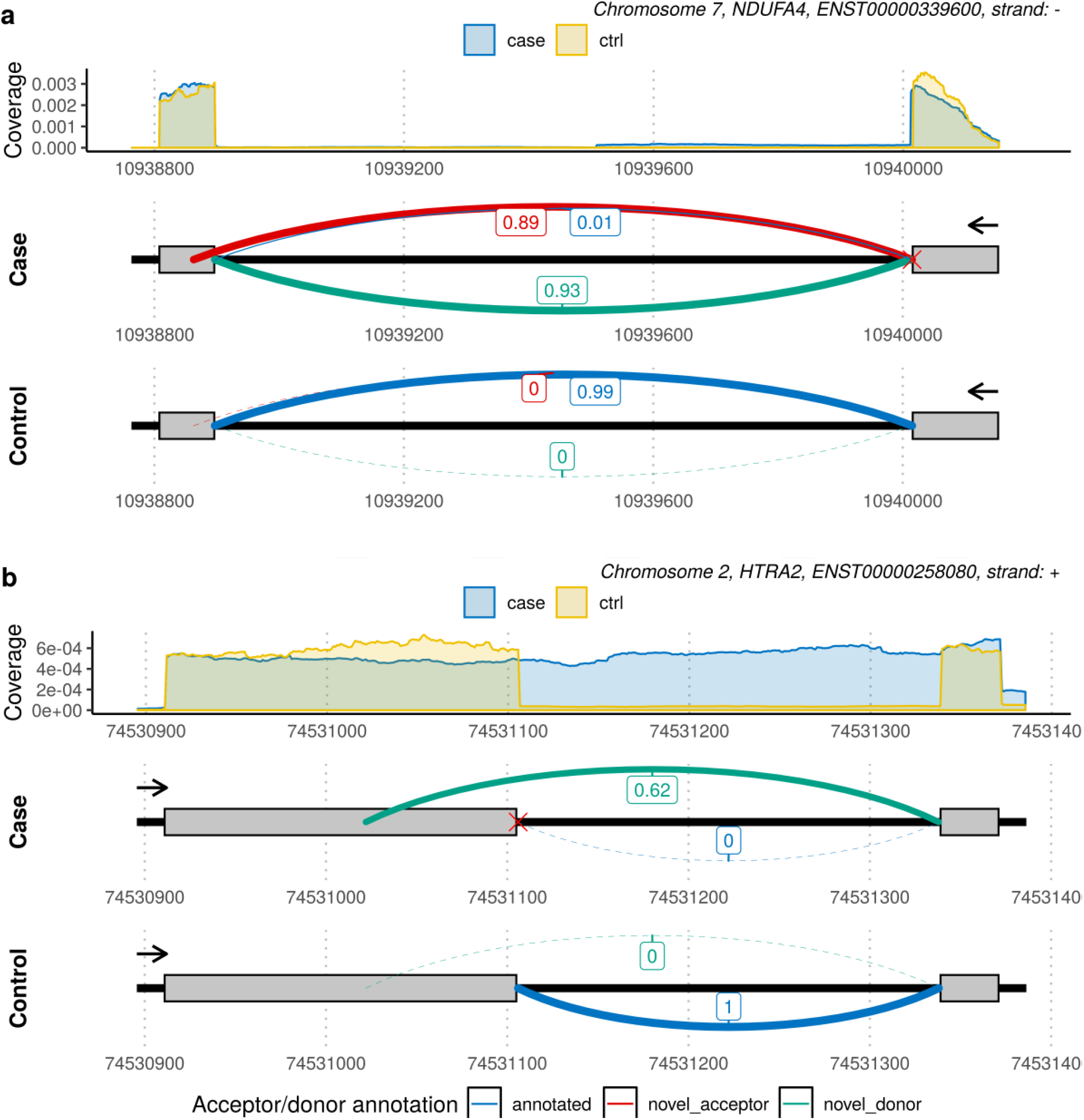
Pathogenic splicing is characterized by disruptions to junctions and coverage. Sashimi plots are split into two panels; the top representing the coverage and the bottom the junctions as well as gene body. Junctions are labelled with their counts and colored with respect to their annotation. The red cross represents the known pathogenic variant. The arrow represents the direction of transcription. **a)** In NDUFA4, the pathogenic splicing event can be observed through the appearance of 2 novel junctions; a novel acceptor (red) and a novel donor (green) junction, which are never found in control samples. Additionally, there is an almost complete loss of an annotated junction (blue), which is always present in control samples. Abnormalities can also be detected in the coverage data across introns associated with the aberrant junctions. First, we can see a slight shift in the right-most exon boundary, which matches the donor site that is represented by the novel donor junction. We can also see a lowly expressed, longer extension of the exon boundary, which is corroborated by the annotated junction that has a normalized count of 0.01 in the patient. **b)** Previous studies have demonstrated that the pathogenic splicing in HTRA2 causes an intron retention event. From the RNA-seq data, we can observe that this is consistent with the loss of an annotated junction (blue) as well as a significant increase in coverage across the intron that is retained. Unexpectedly, there is also an appearance of a novel donor junction (green).

## Methods

### Obtaining the set of OMIM-morbid and gene panel genes

The full set of Online Mendelian Inheritance in Man (OMIM) morbid genes were obtained using the biomaRt R package (v2.40.5), with gene symbols taken from the Ensembl v100 database. The Genomics England panels for neuromuscular disorders (v5.9) and mitochondrial disorders (v2.12) were downloaded from the PanelApp website (https://panelapp.genomicsengland.co.uk/panels/). Only “green” level genes with a high degree of confidence of association with disease were retained for downstream analyses.

### Patient samples

RNA-sequencing was performed on a total of 55 individuals. 16 of these were genetically diagnosed Mendelian disease patients with known pathogenic splicing variants detailed in Supplementary Table 1. The remaining 39 samples were used as controls. Variant types were classified by their proximity to annotated acceptor or donor splice sites. Those within 10bp of an acceptor or donor site were classified as “acceptor” or “donor” variants respectively, whilst those further than 10bp away were termed deep intronic.

### Fibroblast culture and RNA extraction

Fibroblast cell lines cultured in Dulbecco’s Modified Eagle Medium (DMEM) supplemented with 10% Fetal Bovine Serum and 0.05 g/ml uridine. Fibroblasts were harvested by first detaching cells using TrypLE Enzyme, followed by washing with Dulbecco’s Phosphate Buffered Saline (DPBS) prior to storage at −80°C. Total RNA was extracted from fibroblast pellets following the manufacturer’s protocol. In order to assess RNA quality, RNA integrity numbers (RIN) were measured using Agilent Technologies 2100 Bioanalyzer or Agilent 4200 Tapestation with all RIN values found to exceed 8.0.

### RNA-sequencing, alignment and quality control of patient samples

We prepared libraries for sequencing using the Illumina TruSeq Stranded mRNA Library Prep kit by loading 50 ng of total RNA into the initial reaction; fragmentation and PCR steps were undertaken as per the manufacturer’s instructions. Final library concentrations were determined using Qubit 2.0 fluorometer and pooled to a normalized input library. Pools were sequenced using the Illumina NovaSeq 6000 Sequencing system to generate 150 bp paired-end reads with an average read depth of ~100 million reads per sample. Pre-alignment quality control including adapter trimming, read filtering and base correction were performed using fastp, an all-in-one FASTQ preprocessor (v0.20.0)^25^. Reads were aligned using STAR 2-pass (v2.7.0) to the hg38 build of the reference human genome (hg38) using gene annotation from Ensembl v97^26^. Novel junctions discovered in the 1st pass alignment were used as input to the 2nd pass to improve the sensitivity of junction detection. Reads were required to uniquely map to only a single position in the genome. The minimum required overhang length of an annotated and unannotated junction was set to be 3 and 8 base pairs, respectively. The output BAM files underwent post-alignment QC using RSeQC (v2.6.4), with all samples passing quality control after manual assessment^27^.

### Control RNA-seq data

*dasper* analysis was conducted with two sets of controls samples; 504 GTEx (v8) fibroblast samples or a set of 55 in-house samples (including the 16 patients). GTEx v8 fibroblast junction and BigWig data was downloaded via the recount3 R package (v1.1.2) and filtered for samples without large CNVs or chromosomal duplications and deletions (SMAFRZE = “RNASEQ”)^28–30^. In-house RNA-seq data in the form of BAM files were converted into the BigWig format using megadepth (v1.08b) for input into *dasper* (v1.1.3)^31^.

In order to investigate the effect of changing the number of control samples used on the detection of pathogenic events, we down-sampled control numbers systematically. For GTEx control samples, 10, 20, 40, 80, 160, 320 up to a maximum of 504 samples were used. For in-house control samples, analysis was performed using 2, 4, 8, 16, 32 up to a maximum of 55 samples. For each size (N) and type of control samples, 5 iterations were executed. For each iteration, we used N randomly selected control samples of the appropriate type as input into the *dasper* pipeline. When using in-house samples, to ensure that we were not including related patients as controls, any controls with pathogenic variants matching the current patient of interest were removed prior to down-sampling.

### LeafCutterMD

STAR outputted junctions were wrangled into a bed format for input into *LeafCutterMD* (v2.7). All 55 in-house samples were used for intron clustering. Introns were clustered matching the settings used on the *LeafCutterMD* documentation, namely requiring at least 50 junctions supporting a cluster and permitting introns of up to 500kb in size. Outlier intron excision analysis was performed on the 16 patient samples using default settings. Outputted p-values were standardized to ranks for comparison with the output of *dasper*^22^.

#### dasper

Figure 2a depicts the top-level workflow for *dasper* described in the following section. The inputs for *dasper* (v1.1.3) were junction read files (containing reads mapping with a gapped alignment to the genome) and BigWig files (which store coverage data) for control samples and the case sample of interest. Junction reads were annotated based on: i) whether their start and/or end position precisely overlapped with an annotated exon boundary, and ii) whether that junction read matched an intron definition from existing annotation as defined by Ensembl v97^32^. Using this information together with the strand, junctions were categorised as: annotated, novel acceptor, novel donor, novel combination, novel exon skip, ambiguous gene and unannotated. Annotated junctions were those that matched an existing intron definition. Novel acceptor and novel donor junctions had a single end that overlapped with a known exon boundary. Novel combination, novel exon skip and ambiguous gene junctions had both ends overlapping known exon boundaries, however the resulting introns did not match an existing intron definition as defined within Ensembl v97 (SUPP FIG 1). Novel combination junctions connected to exons associated with multiple transcripts, whilst novel exon skip junctions were only associated with a single transcript. Ambiguous gene junctions were connected exons originating from 2 different genes. Unannotated junctions had neither end overlapping a known exon boundary. Junctions were filtered for those that had at least 5 counts in at least 1 sample, a length between 20-1,000,000 base pairs, did not overlap any ENCODE blacklist regions and were not classified as ambiguous gene or unannotated^30^. For each junction, any other junction that shared an acceptor or donor site with it was obtained to form a junction cluster. In order to normalize the junction counts to enable comparison between samples, the counts for each junction were divided by the total counts associated with its corresponding cluster.

**Figure 2.**
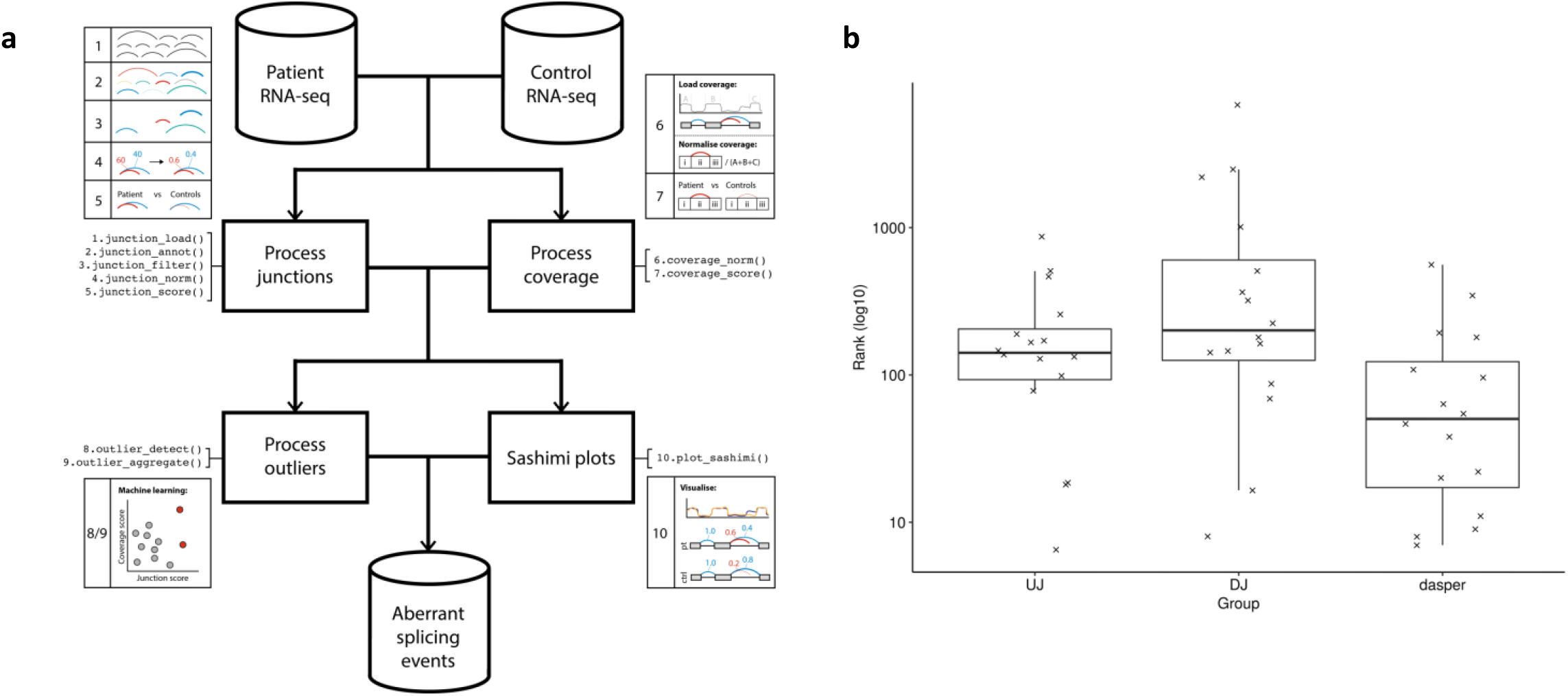
dasper applies an outlier detection method with junction and coverage information as input to detect aberrant splicing events. **a)** dasper takes as input RNA-seq data from a set of cases and controls. Controls can be patient samples or publicly available data, of which dasper includes GTEx data originating from any clinical accessible tissue. Junctions and coverage data are extracted from the RNA-seq and processed. Specifically, this involves normalizing the junction and coverage counts to permit comparison between cases and controls. Then, scoring junctions and coverage by the deviation of their counts from the corresponding count distribution in control samples. These scores are aggregated using an outlier detection model. For each patient, the outputted outlier scores are ranked, generating a list of all splicing events in each patient ranked by their aberrancy. A rank of 1 specifies the most aberrant splicing event in each patient. dasper includes functions to plot sashimi plots to permit manual inspection of candidate splicing events. b) Boxplots displaying the rank of 16 pathogenic splicing events across varying inputs. Each point represents a pathogenic splicing event from one of the 16 patients analyzed. The x-axis shows what information has been used for the ranking, either only up-regulated junctions (UJ), down-regulated junctions (DJ) or the dasper method (UJs, DJs and coverage). The y-axis displays the rank outputted from dasper, with lower ranks specifying splicing events that are predicted to be more aberrant.

For each junction, 3 regions of interest were defined and used to obtain coverage information, namely the intron and the two flanking exons. Exon boundaries were based on exon definitions if the end of a junction overlapped an annotated exon. Otherwise, the putative unannotated exons were presumed to be 20bp in length. Coverage across these 3 regions was loaded from BigWig files. In order to normalize the coverage for comparison between samples, the mean coverage across each of the 3 regions was divided by the total coverage across the exons of the associated gene.

We used z-scores to assess the degree to which junctions and coverage in each patient deviated from the corresponding distribution in controls. For each junction, the coverage z-score with the greatest absolute value across the 3 regions was retained, reducing the number of z-scores per junction from 4 to 2. Junctions were then split into those which had a junction count based z-score above 0 (up-regulated) and below 0 (down-regulated). An isolation forest model was fitted on the up-regulated and the down-regulated junctions separately, using the two z-scores as input. Isolation forests are an ensemble-based outlier detection method, that detect anomalies as those that require shorter paths to isolate^33^. The output of the isolation forest model was an outlier score per junction. Junction-level outlier scores were aggregated to a cluster-level rank in 3 steps. First, clusters that did not contain at least 1 up-regulated and 1 down-regulated junction were omitted. Then, a mean was taken of the up-regulated and down-regulated junction with the greatest outlier scores in each cluster; this formed the cluster-level outlier score. Finally, within each patient, clusters were ranked based on this cluster-level outlier score, with a rank of 1 describing the cluster that had the lowest outlier score and so was predicted to be the most aberrant.

## Results

### Pathogenic splicing events are characterized by abnormalities of annotated junction reads and coverage in associated regions

Previous methods to detect aberrant splicing have often focused on up-regulated novel junctions that are never or very rarely present in controls^15,18^. However, studies have demonstrated that splicing disruptions have complex consequences, which can be difficult to predict from DNA sequence data alone^31^. For this reason, we first explored the consequences of pathogenic splicing variants using RNA-seq data derived from 16 well-characterized and deeply-sequenced patient fibroblast samples. Importantly, this cohort was selected to be heterogenous with respect to disease and variant type [SUPP TABLE 1]. Patient samples were derived from individuals diagnosed with a range of neurological disease, focusing specifically on Mendelian mitochondrial disorders and rare neuromuscular conditions including Ullrich congenital muscular dystrophy [SUPP TABLE 1]. All patients had diagnostically-confirmed splicing variants impacting on acceptor sites, donor sites or located deep within intronic sequence. Detailed inspection using sashimi plots of the resulting sequence data demonstrated that all pathogenic splicing events were characterized by: i) up-regulated novel junction/s (termed UJs), ii) down-regulated annotated junction/s (termed DJs), and iii) changes in coverage within the associated exonic or intronic regions. For example, analysis of RNA-seq data from an individual with a pathogenic donor splice site variant in the gene, *NDUFA4*, confirmed that this variant resulted in the generation of an UJ due to use of an novel donor site 4bp downstream of the canonical splice site^34^ [FIGURE 1a]. However, based on the RNA-seq data we observed additional splicing changes, namely an almost complete absence of an annotated DJ, the appearance of another UJ as well as disruptions in coverage across the first intron [FIGURE 1a]. Similarly, inspection of RNA-seq data derived from an individual with a pathogenic donor splice site variant in the gene *HTRA2* [FIGURE 1b], showed that as well as causing retention of the intron 3 with loss of the canonical splicing event (DJ), there was also a novel UJ caused by use of an novel donor site which was not previously predicted or detected^35^ [FIGURE 1b].

Next, we investigated the relationship between disruptions to junction usage (both UJs and DJ) and abnormalities in sequencing coverage over implied exonic and intronic regions for pathogenic splicing events. This was achieved by calculating corresponding z-scores for each of the four features of interest (UJ, UJ-related coverage, DJ and DJ-related coverage) and based on the distributions of counts and coverage in controls (~50 in-house samples). We found that absolute UJ z-scores were significantly higher than DJ z-scores (median DJ: −15.21, UJ: 27.82, p-value: 0.043) and that both types of junction z-scores tended to be higher than coverage z-scores. Furthermore, we found that the correlation between junction and coverage z-scores was low (Pearson r = −0.1), suggesting that they contained distinct information. Similarly, UJ and DJ z-scores, though negatively correlated (Pearson r: −0.58), could be independently informative for detecting pathogenic splicing events [SUPP FIGURE 2]. Thus, taken together our analysis suggested that pathogenic splicing events were characterized by abnormalities in UJs, DJs and nearby coverage, and that all these features could be informative.

### Development of a clinically accessible, machine-learning pathogenic splicing detection method

Informed by our characterization of pathogenic splicing, we next sought to improve on existing approaches for the identification of aberrant splicing through development of a new tool, *dasper*. Given that we found that pathogenic splicing variants generate both DJs and UJs within a junction cluster, *dasper* explicitly requires each splicing event to have both features, reducing the search space for pathogenic events [FIGURE 2a]. Furthermore, *dasper* incorporates coverage information alongside junction counts to better inform the detection of pathogenic splicing events. These key improvements are embedded within the *dasper* workflow, which begins with the input of patient RNA-seq data, and a set of user-defined RNA-seq control samples. The formats of the files required for *dasper* are standard tabular junction data and BigWigs (Methods: dasper). This enables easy access to large publicly available control data sets through resources such as recount2^29^ and recount3^30^. Leveraging this advantage, *dasper* includes the functionality to download GTEx control data for all clinically-accessible tissues (fibroblasts, skeletal muscle, whole blood, adipose tissue, lymphocytes), permitting the running of *dasper* with only a single patient RNA-seq analysis sourced from any of these sample types. Within *dasper* the user can then load, locally normalize and score junctions and coverage counts in patients based on their deviation from the set of controls (Methods: dasper). After generating junction and coverage-related features, *dasper* applies an outlier detection method, namely an isolation forest, to aggregate junction and coverage scores in a single metric describing the aberrancy of each splicing event^33^. Notably, *dasper* permits easy interchange of the statistical models used to score junctions and coverage as well as the addition of other features, enabling further optimisation of the pipeline in future. Finally, the output of *dasper* is a ranked list of splicing events within each patient sample such that a rank of 1 represents the splicing event predicted to be most pathogenic [FIGURE 2a]. This is complemented by functions that enable visualisation of junctions and coverage of cases and controls in the form of sashimi plots to aid interpretation [FIGURE 1, FIGURE 2a].

We assessed the utility of *dasper* and specifically the value of pairing the use of UJs and DJs, and incorporating coverage information to detect pathogenic splicing events, we compared the ranking of junctions generated on the basis of: i) UJs alone, ii) DJs alone, and *dasper* (UJs, DJs and coverage). This analysis demonstrated that *dasper* ranks pathogenic splicing events on average in the top 34 most aberrant, whilst use of only UJ or DJ information results in average similar ranks of 142 and 202 respectively [FIGURE 2b]. Overall, the use of information originating from both DJs and UJs, alongside the incorporation of coverage in *dasper* improves the detection of pathogenic splicing events.

### dasper outperforms other methods used to detect pathogenic splicing

Next, we evaluated and compared *dasper*’s performance in comparison to existing, commonly used approaches for pathogenic splicing detection, such as LeafCutterMD and z-score^21^. In order to enable comparison between tools, we converted LeafCutterMD p-values and z-scores to ranks such that the lowest p-value or highest absolute z-score was assigned a rank of 1. Based on the analysis of patient-derived fibroblast samples [SUPP TABLE 1], we found that the rankings for pathogenic splicing events produced by *dasper* were significantly lower than those generated by other methods (t-test p-value: vs *LeafCutterMD* 0.013; vs z-score: 0.0003) [FIGURE 3a]. However, given that pathogenic splicing can vary in its difficulty of detection, we also investigated the performance of *dasper* across different variant types both alone and relative to existing methods. We found that while *dasper* detected variants at donor versus acceptor sites with similar accuracy (wilcoxon p-value: 0.482), pathogenic events caused by deep intronic variants received significantly higher ranks, indicating that they were more difficult to detect (wilcoxon p-value: 0.013) [FIGURE 3c]. Nonetheless, we noted that when compared to existing methodologies, *dasper* still had a significantly better performance across all variant types rather than being specific to certain classes. This suggests that *dasper* improves on the detection of pathogenic splicing compared to existing methods, including pathogenic splicing variants that are not in canonical splice sites and which can be difficult to detect.

**Figure 3.**
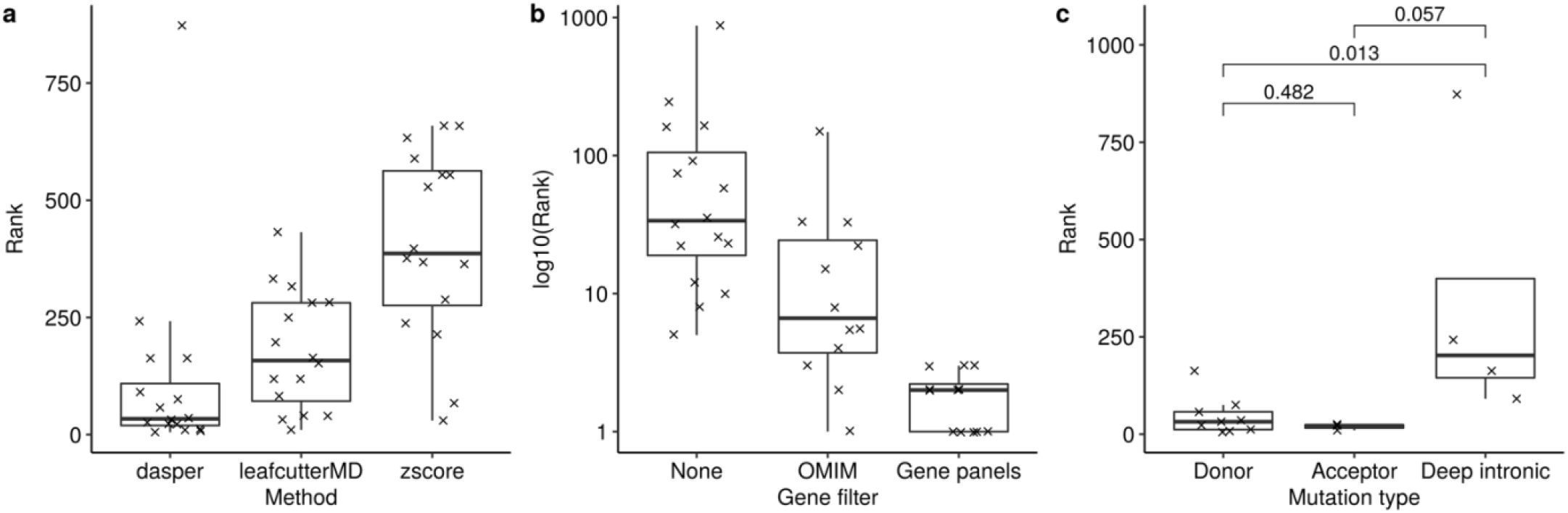
dasper is able to detect pathogenic splicing events more effectively than existing methods and in the top 5 most aberrant splicing events. **a)** Comparison of different methods used to detect aberrant splicing. dasper ranks pathogenic splicing lower or more aberrant than existing LeafCutterMD or z-score approaches. The y-axis represents the rank of pathogenic splicing events, whilst the x-axis specifies the method used**. b)** Ranking pathogenic events across different gene filters. The x-axis details the sets of gene sets that have been used for filtering; either no filter, splicing events connected to OMIM-morbid genes or splicing events associated with gene panels. After applying the OMIM-morbid and gene panel filter, pathogenic splicing events are ranked on average in the top 10 and top 5 most aberrant splicing events respectively. c) Pathogenic splicing events resulting from deep intronic variants are ranked higher than acceptor or donor variants, suggesting that they are more difficult to detect.

We recognized that the utility of *dasper* in diagnostic settings depends not only on how it compares to existing tools but on its performance in clinically-relevant contexts. To investigate this, we measured the absolute ranking of pathogenic splicing events using *dasper*. We found that pathogenic splicing events were ranked on average in the top 40 (median: 33.750) most aberrant events in each patient [FIGURE 3b], but note that these ranks were obtained without any gene, variant or phenotypic level filters. Given that in diagnostic settings only genetic variants in known disease-associated genes would be considered, we re-calculated rankings after filtering for splicing events that were connected to genes within the OMIM-morbid gene set or the appropriate Genomics England panels (see detailed methods). After filtering for OMIM-morbid genes, we found that *dasper* was able to rank pathogenic splicing events within the top 10 most aberrant in each patient (median: 6.750) [FIGURE 3b]. The more stringent gene panel-based filtering, which not only assumes the gene has to be known to cause disease but is also linked to the patient phenotype, further reduced rankings such that pathogenic events were within the top 5 most aberrant on average (median: 2.5) [FIGURE 3b]. In summary, *dasper* is able to rank pathogenic splicing events such that they would be identifiable with only minimal manual curation.

### dasper is able to leverage publicly available control data effectively

While there is increasing evidence to show that paired patient-derived transcriptomic data can increase the diagnostic yield of WES/WGS, there remain significant barriers to implementing this approach in clinical settings. One such hurdle is the generation or identification of suitable control data. In the previous analyses, we have used ~50 in-house sequenced RNA samples as controls. We are aware that sequencing this number of RNA-seq samples would incur a substantial resource burden on diagnostic labs, which may not be feasible in practice. To address this issue, we assessed the performance of *dasper* when using publicly available GTEx v8 data originating from 504 fibroblasts, matching the tissue of origin of patient-derived RNA-seq data in this study^36^. We found that, on average, using in-house samples resulted in more accurate calling of pathogenic splicing events, when compared to the use of GTEx samples as controls. The improvement in ranking of pathogenic events when using in-house controls was observed in 14/16 patients analysed. This pattern of improvement remained true following filtering for pathogenic splicing events within known disease genes (median no filter GTEx: 90, no filter in-house: 34) [FIGURE 4a, SUPP FIGURE 2]. However, this analysis also demonstrated that the absolute ranking when using publicly available controls may be sufficient to be useful when applied in a more clinically-relevant manner. After limiting splicing events to only those connected to genes already implicated in genetic disease as defined in OMIM, and using GTEx controls, *dasper* was still able to rank true pathogenic splicing events in the top 25 most abnormal events (median: 24.5). Overall, this suggests that while technical variability between patient and controls samples reduces our ability to detect pathogenic splicing events, publicly available control data is a viable alternative to costly, time-consuming in-house data generation.

**Figure 4.**
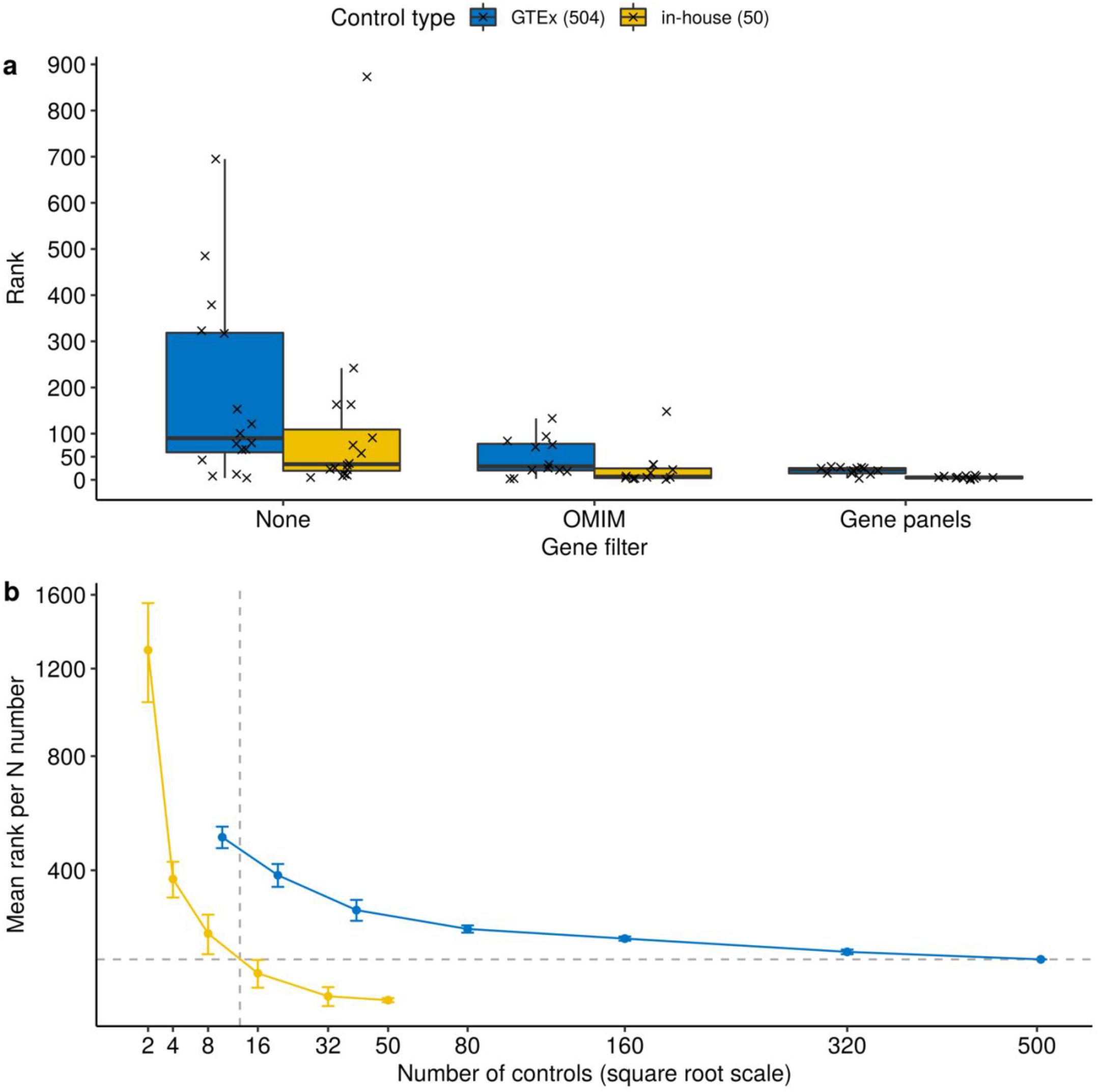
dasper is able to leverage publicly available and in-house controls effectively. **a)** Rank of pathogenic splicing events across varying gene filters and control types. The colour of boxplot represents the control type used, either 504 GTEx v8 samples (blue) or 50 in-house sequenced samples (yellow). In general, in-house samples are able to detect pathogenic splicing events easier than GTEx samples. However, after applying a gene panel filter, pathogenic splicing events are detected in the top 10 splicing events for either control type. **b)** Comparison of the performance of GTEx and in-house control data for detecting pathogenic splicing events. The x-axis describes the number of controls used. The colour of the points and lines describes which control type is used, namely up to 504 GTEx fibroblasts or up to 50 in-house samples. At each N of controls analysed, the mean and standard deviation of the rank of the 16 pathogenic events for the 5 sets of randomly down-sampled controls is plotted.

Next, we explored the relationship between control sample number and the power to detect pathogenic splicing events, a significant concern for implementation in a diagnostic setting whether in-house or external control data is being used. To investigate this, we applied *dasper* while randomly down-sampling the number of control samples used, analysing GTEx and in-house control data separately. As would be expected, we found that an increase in the number of controls considerably improves the detection of pathogenic splicing events using either GTEx or in-house control data [FIGURE 4b]. Notably, while the rate of improvement in pathogenic splicing detection greatly diminishes with increasing control number suggesting a diminishing return, it does not appear to plateau at the maximum number of available samples for either control type. This analysis would suggest that further increases in the quantity of publicly available control samples could compensate against the technical differences between patient and control sample sets. Notably, we found that the ranking when using 504 GTEx controls matched the performance of using between 8 and 16 in-house samples [FIGURE 4b]. In summary, it is likely an increase in sample number would improve the detection of pathogenic events for both control types.

## Discussion

In this study, we present *dasper*, a user-friendly R/Bioconductor package used to detect aberrant splicing events from RNA-seq data. Here, we used a cohort of 16 patients with known pathogenic splicing variants to inform our development of *dasper* and demonstrate its utility. Uniquely, *dasper* pairs information from DJs with UJs as well as incorporating coverage changes across a gene to improve the detection of pathogenic splicing events. As a result, *dasper* outperformed the existing approaches, *LeafCutterMD* and z-score. Furthermore, *dasper* was able to rank pathogenic splicing events in the top 10 most aberrant after OMIM-morbid gene filtering. Designed with clinical accessibility and interpretation in mind, *dasper* uses standard RNA-seq data formats as input granting users flexibility to incorporate publicly available datasets as controls. Moreover, we demonstrated that *dasper* was able to leverage publicly available GTEx data effectively. Finally, *dasper* includes sashimi plot functionality to aid the manual inspection of candidate splicing events^28,29,37^.

To the best of our knowledge, this is the first study to explore the impact of splicing variant subtypes and control sample selection. We demonstrate that *dasper* outperforms existing tools (*LeafCutterMD* and z-score) on splicing variants located not only at donor and acceptor sites, but also at deep intronic splicing variants, which are most challenging for all existing methods. Furthermore, our analyses highlighted that selection of control sample type and number greatly impacts on the power of pathogenic splicing detection. In particular, we compare two approaches to obtain control data; either the use of publicly available GTEx RNA-seq data or in-house sequencing data. While we find that the use of in-house samples improves the performance of *dasper*, presumably because of a reduction in the technical differences between patient and control data, this approach is associated with increased costs and reduced flexibility, creating barriers to the use of RNA-seq pipelines in diagnostic laboratories. In contrast, the use of publicly available datasets has minimal associated costs and is highly flexible, both in terms of the clinically-accessible tissue samples that can be analysed for a given patient, and the batching of samples which significantly affects turnaround times for laboratory results. Although *dasper* performed better when using in-house samples, GTEx samples still enabled pathogenic splicing events to be detected, on average, in the top 25 most aberrant after applying an OMIM-morbid filter alone. This ranking was equivalent to using 8-16 in-house samples suggesting that use of publicly available data is a viable, cost-effective alternative for the detection of pathogenic splicing. In this context, it is worth noting that public RNA-seq datasets are progressively increasing in size. In fact, for tissues such as blood, public datasets used collectively could provide RNA-seq profiles for >30,000 unrelated individuals which could be meta-analysed as elegantly demonstrated by the eQTLgen consortium^38^. Additionally, over 70,000 and 300,000 human RNA-seq samples are in recount2 and recount3 respectively^28–30^. Publicly available control data sets at this scale have the potential to both surpass the importance of large in-house control data sets, and permit different centers to use identical computational protocols for diagnoses that enable the standardization of pathogenic splicing identification across laboratories.

Together, through leveraging in-house or publicly available datasets effectively, we hope *dasper* will make RNA-seq a more affordable, effective and standardized tool for diagnostics and ultimately, lead to an increased rate of genetic diagnosis.

## Supplementary figures

**Supp Figure 1.**
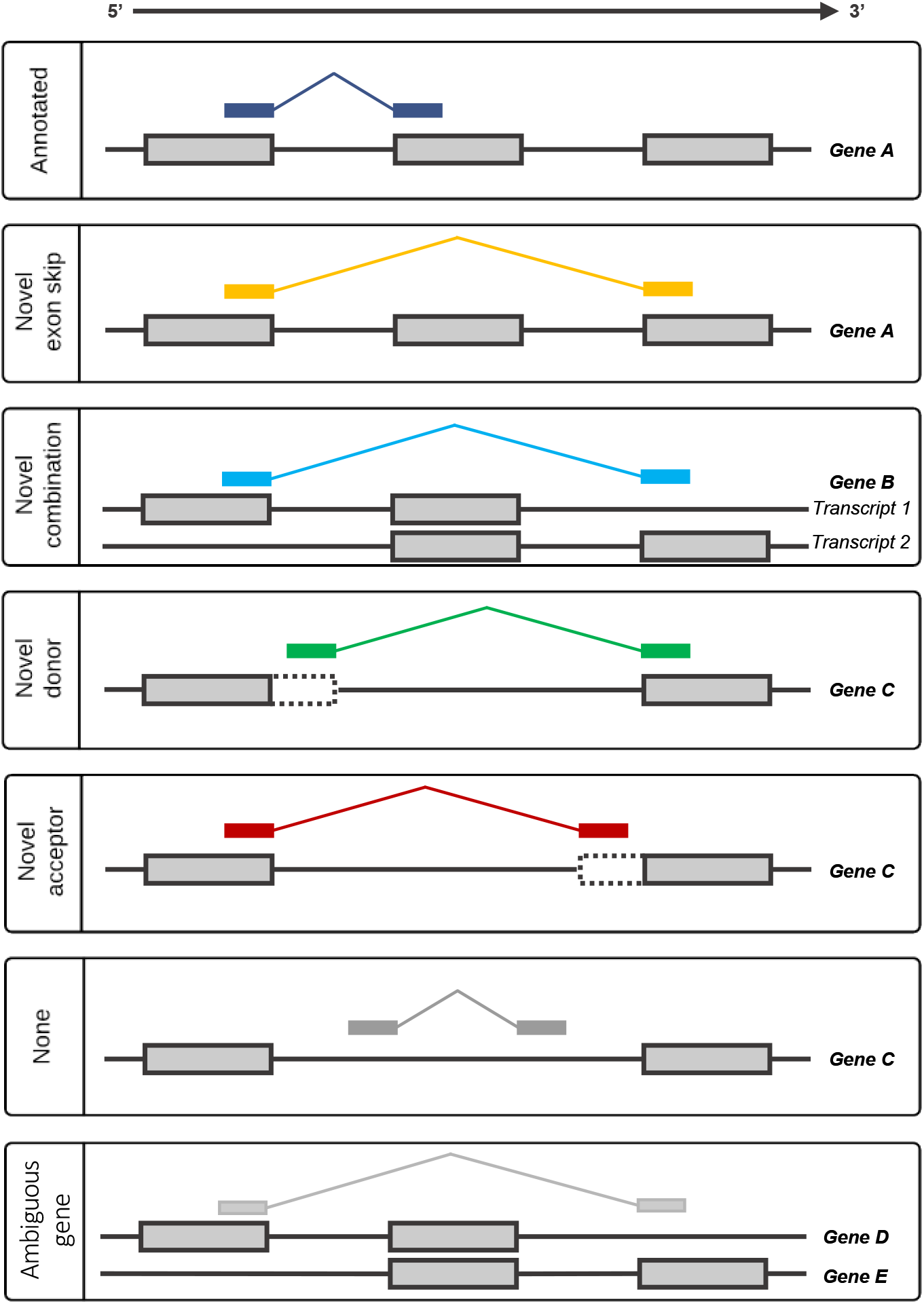
Schematic illustration of the different categories of splicing event. Junction reads used to define Leafcutter introns were annotated based on their relationship to the annotated transcriptome (Ensembl v97). Here, the annotated transcriptome is illustrated by the grey-filled boxes. Annotated junctions have donor and acceptor splice sites that match the boundaries of an existing intron. Likewise, novel exon skip and novel combination junctions have donor and acceptor splice sites that overlap known exon boundaries derived from exons contained within the same transcript, but, they represent introns which are not found in the set of annotated introns. They are distinguished by whether or not their donor and acceptor splice sites overlap exons derived from the same transcript. Novel donors and novel acceptors are junctions where only one end (3’ or 5’, respectively) matches the boundary of a known exon. All novel events are considered partially annotated. Unannotated junctions (“None”) have neither end overlapping a known exon. Ambiguous gene junctions are have either end overlapping different genes.

**Supp Figure 2.**
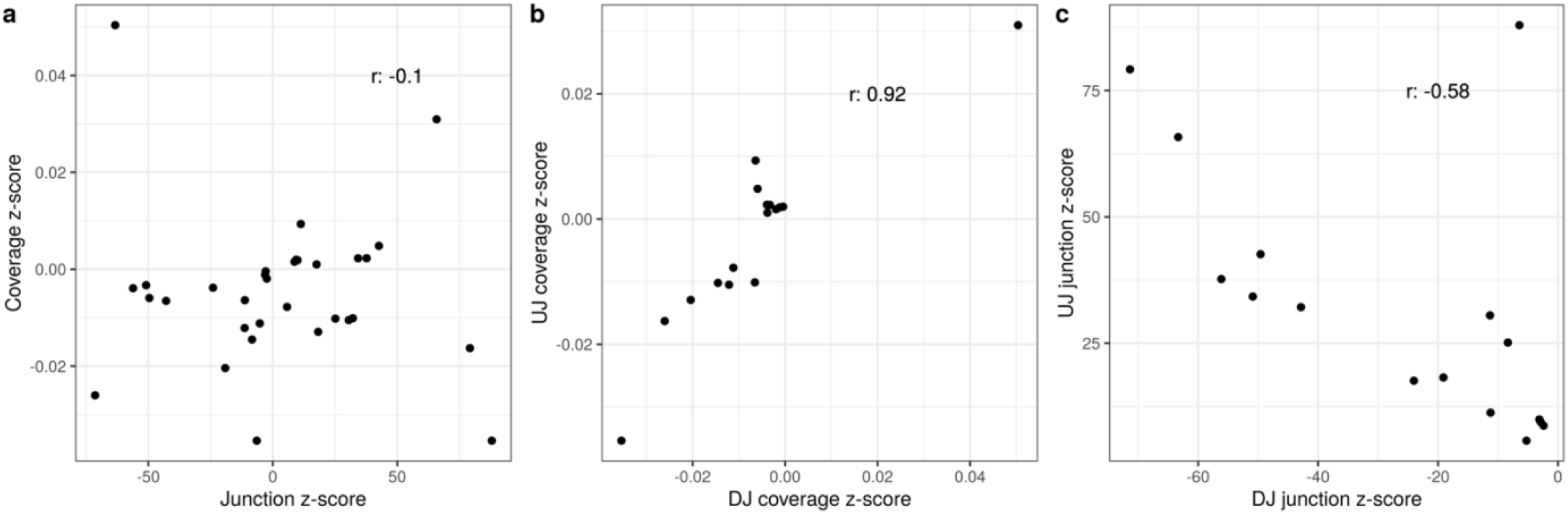
Correlation z-scores that were used as input into dasper. Each point represents a particular patient sample. A Pearson correlation was run with the r values specified on the plot. **a)** Correlation between coverage and junction z-score. **b)** Correlation between the coverage z-scores for up and down regulated junctions. **c)** Correlation between junction z-scores for up and down regulated junctions.

## Acknowledgements

M.R. was supported by the UK Medical Research Council (MRC), and the award of a Tenure Track Clinician Scientist Fellowship (MR/N008324/1). R.H.R. was supported through the award of a Leonard Wolfson Doctoral Training Fellowship in Neurodegeneration. The University College London Hospitals/University College London Queen Square Institute of Neurology sequencing facility receives a proportion of funding from the Department of Health’s National Institute for Health Research Biomedical Research Centres funding scheme. The clinical and diagnostic ‘Rare Mitochondrial Disorders’ Service in London is funded by the UK National Health Service (NHS) Highly Specialised Commissioners. R.D.S.P. is supported by a Medical Research Council (MRC) Clinician Scientist Fellowship (MR/S002065/1) and receives funding from a Medical Research Council (MRC) strategic award to establish an International Centre for Genomic Medicine in Neuromuscular Diseases (ICGNMD) (MR/S005021/1). R.M. and R.W.T. are supported by the Wellcome Centre for Mitochondrial Research (203105/Z/16/Z), the Medical Research Council (MRC) International Centre for Genomic Medicine in Neuromuscular Disease (MR/S005021/1), the Mitochondrial Disease Patient Cohort (UK) (G0800674), the Lily Foundation and the UK NHS Specialised Commissioners who fund the “Rare Mitochondrial Disorders of Adults and Children” Service in Newcastle upon Tyne. J.J.C. was supported by a Barbour Foundation PhD studentship and an EMBO Short Term Fellowship. The MRC Centre for Neuromuscular Diseases Biobank, for providing patients’ samples. H.Z and F.M were supported by the Muscular Dystrophy UK, and the National Institute of Health Research (NIHR) Biomedical Research Centre at Great Ormond Street Hospital and UCL.

## Author contributions

D.Z. and M.R. conceived and designed the study. D.Z. analyzed the data, generated figures, and together with M.R. wrote the first draft of the manuscript. L.C.T., R.H.R., and M.R. helped guide and troubleshoot analyses. L.C.T. and. D.Z. contributed to submitting dasper as a R package to Bioconductor. S.A. I.A.B, J.J.C, H.H, R.M., F.M, M.O., J.P. M.S., R.D.S.P, R.W.T, H.Z. and C.D collected and facilitated the obtaining of patient samples. R.H.R., S.G, E.G, S.S, J.B, S.A. J.J.C, H.H, F.M, M.O., R.D.S.P, R.W.T, H.Z. and M.R. contributed to the critical analysis of the manuscript.

